# Neural epidermal growth factor-like 1 protein variant increases survival and modulates the inflammatory and immune responses in human ACE-2 transgenic mice infected with SARS-CoV-2

**DOI:** 10.1101/2021.02.08.430254

**Authors:** Roopa Biswas, Shannon Eaker, Dharmendra Kumar Soni, Swagata Kar, Denae LoBato, Cymbeline Culiat

**Author notes:** Corresponding Authors Roopa Biswas, Department of Anatomy, Physiology and Genetics, Room B4024, Uniformed Services University of the Health Sciences, 4301 Jones Bridge Road, Bethesda, Maryland 20814. Email address; Cymbeline Culiat, NellOne Therapeutics Inc., 11020 Solway School Road, Suite 101, Knoxville, TN 37931. Contributed equally to this manuscript (Roopa Biswas, Cymbeline Culiat).

## Abstract

Coronavirus disease 2019 (COVID-19) is a viral illness caused by the severe acute respiratory syndrome coronavirus 2 (SARS-CoV-2) and is a worsening global pandemic. COVID-19 has caused at least 1.7 million deaths worldwide and over 300,000 in the United States. Recently, two promising vaccines are being administered in several countries. However, there remains an urgent need for a therapeutic treatment for COVID-19 patients with severe respiratory damage that can lead to intensive care, prolonged hospitalization, or mortality. Moreover, an increasing population of patients manifest lingering disabling symptoms (called Long Haulers). Here, we tested the efficacy of a recombinant neural epidermal growth factor like 1 protein variant (NELL1-NV1) in a COVID-19 mouse model, transgenic mice expressing the human angiotensin I-converting enzyme 2 (ACE2) receptor (tg-mice hACE2) infected with SARS-CoV-2. The administration of NELL1-NV1 to SARS-CoV-2-infected tg-mice hACE2 significantly improved clinical health score and increased survival. Analyses of bronchoalveolar (BAL) fluid demonstrated decreased levels of several cytokines and chemokines (IFN-γ, IL-10, IL-12 p70, CXCL-10/IP-10, MIG and Rantes), in NV1-treated treated mice compared to controls. Cytokines including IL-1α, IL-9, IL-6, LIX/CXCL5, KC/CXCL1, MIP-2/CXCL2, MIP-1α/CCL3, and G-CSF, critical to immune responses such as neutrophil recruitment, viral clearance and vascularization, were increased compared to controls. Our data suggest the potential of NELL1-NV1-based therapy to mitigate the cytokine storm, modulate the abnormal immune response and repair respiratory tissue damage in COVID-19 patients.

## INTRODUCTION

The coronavirus disease 2019 (COVID-19) is a viral illness caused by the severe acute respiratory syndrome coronavirus 2 (SARS-CoV-2) that has spread rapidly in a global pandemic affecting more than 200 countries (https://coronavirus.jhu.edu/map.html; Johns Hopkins University Center for System Science and Engineering COVID-19 Dashboard).^1^ In nearly a year it has caused 1.7 million deaths worldwide and over 320,000 in the United States (World Health Organization Coronavirus Disease COVID-19 Dashboard; https://covid19.who.int/). The disease spectrum ranges from a mild respiratory illness to a severe disease requiring hospitalization and intensive care.^1^ Severe disease is marked by an aberrant immune response often triggering hyperinflammation or a “cytokine storm” that damages lung tissue, compromises oxygen exchange, and can lead to acute respiratory distress syndrome (ARDS) with hypoxia, multiple organ failure, and death in approximately 7% of patients in the U.S.^1-3^ The standard-of-care for this fragile patient population is limited to supportive care (*e*.*g*. oxygen therapy, mechanical ventilation), prolonged stay in an intensive care unit, and the administration of repurposed existing drugs (*e*.*g*. antiviral agents, anti-inflammatory agents).^1^ Progression to ARDS is linked to ∼80% of COVID-19 fatalities, and survivors have significant defects in lung structure and function.^2,3^ Patients who survive severe COVID-19 can manifest lung scarring, fibrosis, and a reduction of lung function even in previously healthy individuals without known risk factors.^4-7^ COVID-19 cases are more prevalent among high-risk patient populations such as the elderly and those with pre-existing medical conditions like obesity, hypertension and diabetes.^1^ Patients with diabetes comprise ∼30% of COVID-19 cases in the U.S., with hospitalizations and deaths 6X and 12X higher, respectively.^8.9^ An increasing new patient population called the “long haulers” consists of individuals who were not serious enough to be hospitalized but suffer lingering symptoms after several months, and is estimated by the CDC to be 20-30% of confirmed COVID19-positive persons.^10-12^ Several studies have also revealed a strong association of the male sex being a risk factor for increased disease severity and mortality (nearly twice the rate).^13,14^

It is thought that the severity of COVID-19 is brought about by aberrant immune responses to the SARS-CoV-2 infection which are highly variable among patient populations, but are often characterized by an aggressive response by inflammatory processes reminiscent of a cytokine release syndrome or “CRS”, mobilization of an abnormal cellular milieu, and the triggering of autoimmune pathways.^15-26^ The immunology (innate and adaptive mechanisms) governing the body’s response to COVID-19 disease process is under intense scrutiny because of the implications to prognosis determination, disease severity, vaccine development and distribution, and therapeutic interventions.^15-21^

High mortality in COVID-19 patients is in part due to acute respiratory distress syndrome (ARDS) post-infection. Researchers from the Cell Therapy field are using previous experiences dealing with CRS, seen in patients receiving T-cell therapies (such as Chimeric Antigen Receptor T-cells/CAR-T), where a similar hyperimmune response is seen.^22-23^ CRS in CAR-T therapy has symptoms (also similar to COVID-19 such as fever, respiratory distress, tachycardia) related to elevated levels of cytokines and chemokines, specifically IL-6 and INF-γ. Current drugs in clinical trials for CRS are now shifting from cell therapy applications to COVID-19 treatments in hopes of calming the post-infection cytokine storm. Treatments for cell therapy-related CRS include the IL-6 inhibitor Tocilizumab, while similar IL-6 inhibitors such as Siltuximab and Sarilumab and Janus kinase (JAK) inhibitors are currently being evaluated for COVID-19. However, treatments with these drugs may come with side effects, such as the reduction of IL-10 which could be needed for viral clearance.^24^ In addition, as targeted therapies against specific cytokines, such as inhibitors against IL-6 (Roche, Sanofi, Regeneron) and GM-CSF (GlaxoSmithKline), are in clinical trials or approved for diseases such as arthritis, these groups are looking to repurpose these drugs for COVID-19. However, as data show that cytokine expression is inconsistent in different global clinical reports and might only benefit a smaller portion of the affected population (i.e. ARDS patients without any increases in IL-6 or IL1β cytokine levels), a more holistic approach should be taken for current and future coronavirus infections such as COVID-19.^20,25,26^ It is critical to understand and treat the immunological response in respiratory viral infections in its entirety, aiming strategies at addressing cytokine signaling to significantly reduce hyperinflammation in patients with severe COVID-19.^27,28^ However, to date, mostly cytokine-specific targeted drugs are being explored. The use of Vitamin D for suppressing the hyperimmune response has been speculated, possibly involving C-Reactive Protein (CRP) regulation.^28^ As current data suggests that both the innate and adaptive immune systems are involved in COVID-19, such targeted therapeutics should be balanced with antiviral support and overall T-cell homeostasis for clinical improvements and viral clearance. Research into growth factors that regulate and balance the immune system response are needed.^20, 21^

NELL1 is a naturally occurring secreted signaling molecule that mediates tissue growth and maturation during fetal development and stimulates healing in skin, muscle, bone, heart and cartilage after acute injury and systemic diseases such as osteoporosis and osteoarthritis.^29-38^ It is essential during the development of epithelial linings of esophagus, gastrointestinal tract and respiratory system.^39-41^ Fifteen years of basic science research on NELL1 biology demonstrate that exogenous NELL1 has strong anti-inflammatory and pro-tissue healing effects in the skin, muscle, cartilage, and bone (**Table 1**).^31-38, 41-43^ A hallmark of NELL1 activity is the modulation of the pro-inflammatory response in systemic inflammatory diseases and acute tissue injuries.^37,41-43^ In contrast to stem cell therapies currently undergoing clinical testing in COVID-19 patients, NELL1 (1) promotes tissue repair without the associated effects of uncontrolled growth and immune system rejection, and (2) is a recombinant protein that does not rely on complex and laborious stem cell resources. Thus, the lack of existing therapeutics for ARDS, coupled with NELL1’s validated strong tissue healing effects in hyper-inflamed and hypoxic tissue injury environments, support the expansion of its applications to mitigate the cytokine storm and respiratory tissue damage induced by SARS-CoV-2. (**Fig. 1**)

**Table 1.** NELL1 regulates molecules implicated in SARS-CoV-2 lung fibrosis. The activity of NELL1 to modulate the levels and activities of pro-inflammatory molecules in other tissue damage context suggested its therapeutic potential for regulating the hyperinflammation and immune cells responses characterizing COVID-19.

| Table 1: NELL1 regulates molecules implicated in SARS-CoV-2 lung fibrosis |  |  |
| --- | --- | --- |
| Molecule | Lung Damage and Fibrosis studies<br>Refs: 17, 70-73 | NELL1 Studies<br>Refs: 32, 37, 41-43 |
| IL-1 $\beta$ cytokine | Rat lung fibrosis model<br>Human lung tissues with ARDS | Human skin UV damage repair *in vitro*<br>Cartilage inflammation *in vitro*<br>Osteoarthritis *in vivo*<br>Human muscle atrophy rescue *in vitro* * |
| IL- 6 cytokine | Human lung tissues with ARDS | Bone regeneration *in vivo*<br>Cartilage Inflammation *in vitro* |
| IL- 8 cytokine | Human lung tissues with ARDS | Human skin UV damage repair *in vitro* |
| TNF-α cytokine | Rat lung fibrosis model<br>Human lung tissues with ARDS | Bone regeneration *in vivo*<br>Cartilage inflammation *in vitro* |
| NF-$\kappa\beta$ cytokine | Mice SARS-CoV infection | Bone regeneration *in vivo* |
| MMP1<br>matrix metalloproteinase | Human lung tissues with idiopathic<br>pulmonary fibrosis and bleomycin rat model;<br>transcriptomics of human lung tissues | Human skin UV damage repair *in vitro* |
| MMP13<br>matrix metalloproteinase | Human lung tissues with idiopathic<br>pulmonary fibrosis and bleomycin rat model | Cartilage Inflammation *in vitro* |

\*unpublished

**Figure 1.**
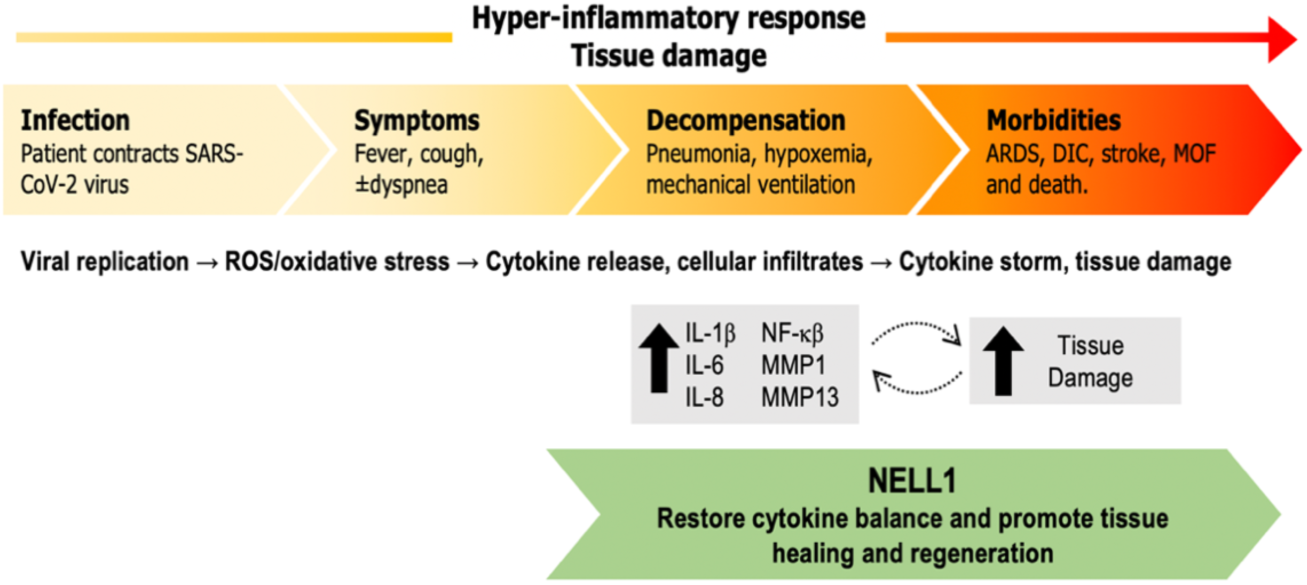
Overview of COVID-19 disease progression and the therapeutic potential of NELL1. (Top) COVID-19 disease progression. During SARS-CoV-2 infection, the host mounts an immune response, which in some patients, triggers an aberrant pro-inflammatory feedback loop (gray) that compromises both structural tissue integrity and functional oxygen exchange, leading to acute lung injury and acute respiratory.

NV1 is a variant of the endogenous NELL1 with improved properties for commercial manufacturing and soft tissue healing (unpublished). A NELL1 therapeutic has great potential to reduce COVID-19 morbidity in the near term and prevent long-term tissue damage caused by SARS-CoV-2 or respiratory syndromes caused by other viruses (e.g. influenza viruses H1N1 and H3N2). By addressing both the acute and prolonged tissue injury effects of virus-induced damage to the respiratory system, a NELL1-based therapeutic has the potential to improve survival rates and patient quality-of-life, hence reducing the massive economic burden on patients and healthcare systems.

Many animal models in mice, ferrets, hamsters and non-human primates are being developed and used to investigate the basic biology of SARS-CoV-2 infections, transmission, susceptibility and mechanisms of disease.^44-47^ More importantly, they are also being utilized to evaluate the efficacy and safety of potential therapies against COVID-19. The primary limitations of these animal models were: a) the inefficient infection by the human SARS-CoV-2 virus to the ACE2 receptor of the animal model, and b) the failure of the model to develop the severe COVID-19 features exhibited by hospitalized patients in the instances where viral infection was enhanced by genetic manipulation or passaging.^44,45^ For instance, hamsters, ferrets and even non-human primates only developed mild to moderate disease and recovered spontaneously. Recently, several studies demonstrated that transgenic mice expressing the human angiotensin I-converting enzyme 2 (ACE2) receptor driven by the cytokeratin-18 promoter (k18-hACE2-tg mice) are infected by high levels of SARS-CoV-2 (e.g. 2.5 × 10^4^ plaque forming units delivered intranasally) and recapitulates the symptoms of lung disease and pulmonary failure in severe COVID-19 patients.^45-48^ Increased viral loads correlated with rapid weight loss and mortality, deterioration of multiple respiratory parameters, infiltration of the lungs with immune system cells like monocytes, neutrophils and activated T-cells, and high levels of pro-inflammatory cytokines and chemokines. The K18-hACE2-tg mice therefore, presented as an appropriate model to initially test the potential of NELL-NV1 as an immunomodulatory and therapeutic treatment for severe COVID-19.

## MATERIALS AND METHODS

### NELL1-NV1 recombinant protein

Recombinant NELL1-NV1 gene sequence was synthesized and cloned into the Wuxian Express vector and expressed in Chinese Hamster Ovary cells (stable cell line CHO-K1; WuXi Biologics, Hongkong Ltd). The polypeptide (610 amino acids; 90 kDa) was purified by Ni-excel and Ni-NTA and resuspended in phosphate buffered saline (PBS; pH=7.2) at a concentration of 0.54 mg/mL (Lot No. 20190826-7058). The product was analyzed by SDS-PAGE under reducing and non-reducing conditions, and SEC-HPLC. Purity was determined to be at >95% and endotoxin levels at <1 EU/mg.

### Mouse model for COVID-19

The tg-mice hACE2 (strain 034860 B6.Cg-TG(K18-ACE2) 2Primn/J) were obtained from Jackson Laboratory (Bar Harbor, ME, USA). The experiments with live virus challenge were carried out at BIOQUAL Inc. (Rockville, MD, USA) biosafety level 3 (BSL-3) facilities in compliance with local, state, and federal regulations under IACUC protocol #20-083.

### SARS-CoV-2 infection

SARS-CoV-2 strain USA-WA1/2020 (BEI Resources NR-52281, batch #70033175), courtesy of the Centers for Diseases Control and Prevention (CDC), was used for mice infection. The SARS-CoV-2 stock was expanded in Vero E6 cells, and the challenging virus was collected at day 5 of culture when the infection reached 90% cytopathic effect. Sequencing confirmed that the full genome of the virus in the expanded stock was 100% identical to the parent virus sequence listed in the GenBank database (MN985325.1). A plaque-forming assay was conducted in confluent layers of Vero E6 cells to determine the concentration of live virions, measured as plaque-forming units (pfu). tg-mice hACE2 (8-10-weeks-old; N=15, 8 females, 7 males) were anesthetized with ketamine/xylazine and infected via intranasal injection with 2.8 × 10^3^ pfu of SARS-CoV-2. Bodyweight and clinical observations were monitored daily during the experimental period.

### Administration regime for NV1

Following infection, tg-mice hACE2 were administered with two different doses of NV1: 1.25 mg/kg body weight [BW] (N=5) or 2.5 mg/kg BW (N=5) NELL1 in 100 µL PBS via Retro-Orbital (RO) injections on 0 and 3 dpi. Control mice (N=5) were mock treated similarly with sterile PBS only.

### Profiling of Cytokines and Chemokines

The levels of cytokine and chemokines in BAL samples from NV1-treated and control SARS-CoV-2-infected mice were analyzed by Luminex multiplex assays using 36-plex MILLIPLEX® MAP Mouse Cytokine / Chemokine Magnetic Bead kit (EMD Millipore). BAL samples were quickly vortexed for 10s, then centrifuged to pellet and remove debris. Samples were then analyzed according to the manufacturer’s protocol. Briefly, 200 µL of bead mix solution were added to all wells and washed using a Bio-Plex Pro Wash Station (BIORAD). 25 µL of Universal Assay Buffer were added to all wells, then 25 µL of samples/standards/blanks (blanks utilize universal assay buffer) were added to the relevant wells, following a 2h incubation on an orbital shaker (400-500 rpm) at room temperature. Plates were then washed on the magnetic plate washer, and 25 µL of prepared detection antibody were added to all wells following a 30 min incubation, with agitation, at room temperature. Plates were washed again, following 50 µL of provided Streptavidin-PE solution were added to all wells, 30 min incubation with agitation at room temperature. Once complete, the plates were again washed using the magnetic plate washer, inactivation was then performed using a 10% Formalin solution (VWR), using 100µL per well, overnight. Plates were run on a Bio-Plex^®^ 200 System (BIORAD), which was preprogrammed according to kit specifications for optimal signal detection.

### Statistical Analysis

Analyses of percent change in body weight, Kaplan-Meir survival, and change in cytokine expression in NV1-treated compared to control SARS-CoV-2-infected tg-mice hACE2 were analyzed initially using Prism version 8 (GraphPad). The Mixed-effects Model with Toeplitz errors was used to analyze changes in weight by groups over time. Cox proportional hazard model was used to analyze the survival data. The Kruskal Wallis test was employed to determine statistically significant differences (p = 0.05) in cytokine and chemokine levels between NV1-treated and control groups.

### Histopathology Analysis

Lung tissues were collected, at euthanasia, from hACE2 tg mice infected with SARS-CoV-2 and preserved in 10% neutral buffered formalin. The samples were processed using standard histology techniques for serial dehydration in alcohol, embedding in paraffin, sectioning and staining the tissues with Hematoxylin and Eosin and Masson’s Trichrome (Histoserv Inc., Germantown, MD 20874). Representative tissue specimens were examined from treatment groups (PBS N=3; NV1 treatment at 1.25 mg/kg BW dose, N=5; NV1 treatment at 2.5 mg/kg BW, N=4) by a board certified veterinary anatomic pathologist blinded to treatment group. A comparison to reference normal uninfected lung frozen tissue samples that were embedded, sectioned and stained in the same manner as the infected tissues (Jackson Lab Cat. 034860; N=4) was conducted to identify and confirm lesions associated with the SARS-CoV-2 infection. Slides from treatment group animals were compared to the normal tissues and evaluated for the presence of histologic changes including evidence of inflammation, degree of consolidation, and vascular pathology.

## RESULTS

### Improved clinical scores and survival of SARS-CoV-2-infected tg-mice hACE2 after administration of NV1

We examined the efficacy of NV1 treatment on BW and survival of SARS-CoV-2-infected tg-mice hACE2. The clinical symptoms of virus-infected control mice include abrupt loss in body weight at 1 dpi (**Fig. 2A**, black), with ruffled fur, hunched back, and reduced mobility. On average, SARS-CoV-2-infected tg-mice treated with the lower dosage (1.25 mg/kg BW) of NV1 showed a significant increase in BW (p<0.05), peaking to >10% at 3 dpi (**Fig. 2A**). This group includes 2 male and 3 female mice. Interestingly, the 3 female mice exhibited a significant weight gain and scored normal clinically till day 4 dpi and subsequently, developed clinical signs of mild-ruffled, ruffled fur, hunched back and listlessness. SARS-CoV-2-infected tg-hACE2 mice (N=5) treated with the higher dosage (2.5 mg/kg BW) of NV1 protein showed no significant change in BW (**Fig. 2A**). Furthermore, the Kaplan-Meier survival curve (**Fig. 2A)** indicate a 40% survival for the lower dosage of NV1 at 6 dpi compared to a 20% survival observed for the higher dosage. Statistical analysis did not show significant differences of body weight and survival between male and female mice in this small test group. The trend of greater protection observed in the female mice might become more apparent in a larger population of males and females. These results imply that optimizing the administration regime and dosage and examining further gender-based effects might improve the efficacy of NV1 for COVID-19 therapy.

**Figure 2.**
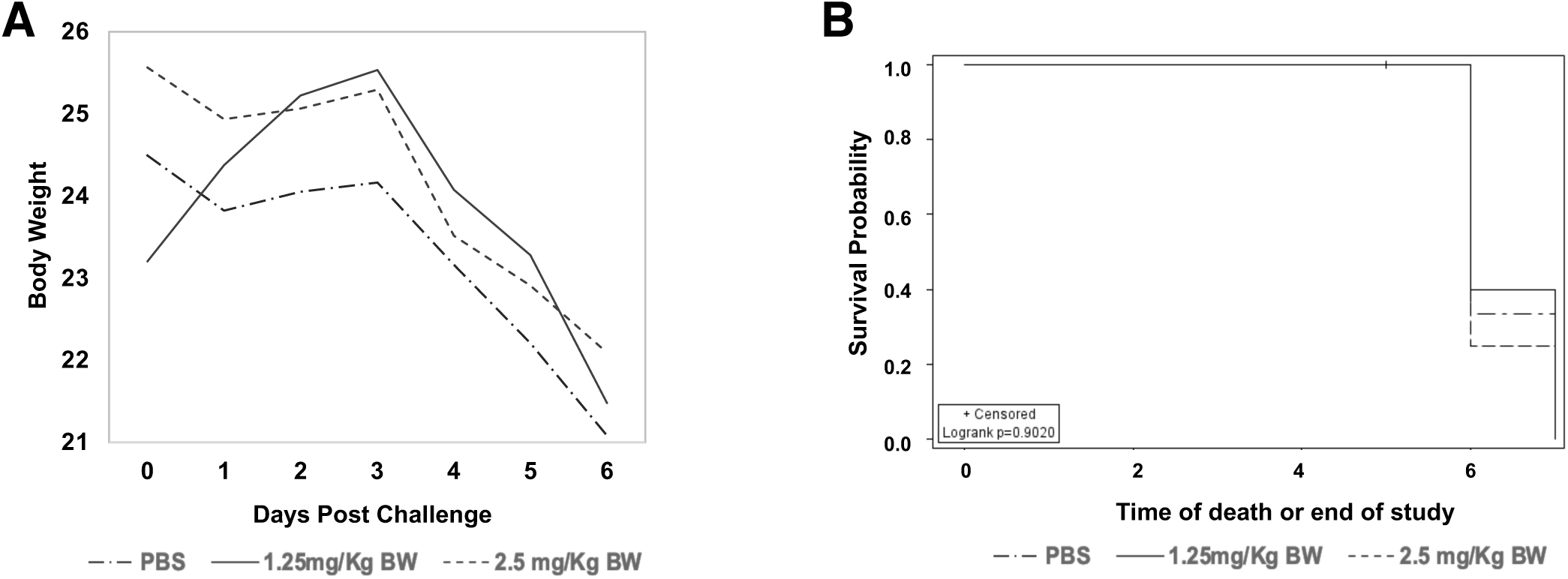
NV1 treatment in SARS-CoV-2 infected tg-mice hACE2. NV1 was administered to tg-mice hACE2 by retro-orbital injection on days 0 and 3 post-infection with SARS-CoV-2. **(A)** The lower dose (1.25 mg/kg BW, N = 5) of NV1 protein induces significant increase in body weight compared to uninfected control mice (p<0.05). The higher dose (2.5 mg/kg BW, N=5) of NELL1 was not as effective. **(B)** The corresponding Kaplan-Meier survival plot indicates a 40% survival with the lower dose of NELL1 and a 20% survival with the higher dose of NELL1.

### NV1 treatment attenuated multiple components of the cytokine storm in SARS-CoV-2-infected-tg-mice hACE2 and maintained high levels of molecules needed for viral immune response

A comprehensive cytokine profile analyses was conducted with undiluted BAL samples collected at euthanasia, by Luminex multiplex assays using 36-plex MILLIPLEX® MAP Mouse Cytokine / Chemokine Magnetic Bead kit (EMD Millipore). The levels of pro-inflammatory markers interferon gamma (IFN-γ), interleukin 10 (IL-10), interleukin 12 p70 (IL-12(p70)), CXC motif chemokine 10 (CXCL-10) also known as interferon gamma-induced 10kDa protein 10 (IP-10), monokine induced by IFN-γ (MIG), and Regulated on Activation, Normal T cell Expressed and Secreted (Rantes or CCL5) were decreased in both NV1 doses (**Fig. 3**). NV1-treated mice displayed high levels of multiple components of the inflammatory cascade (**Fig. 3B**) compared to the control/PBS-treated group. Statistically significant differences in the levels of interleukin 9 (IL-9; p = 0.028), interleukin 1 alpha (IL-1 α; p = 0.02) and lipopolysaccharide-induced CXC chemokine (LIX/CXCL5; p = 0.05) were detected. Although differences in the levels of other molecules in the inflammatory response did not reach statistical significance, strong trends were noted in: interleukin 6 (IL-6; p=0.065), macrophage inflammatory protein 2 (MIP-2 or CXCL2; p= 0.084), keratinocyte chemoattractant (KC or CXCL1; p=0.137), granulocyte colony stimulating factor (G-CSF; p=0.164), macrophage inflammatory proteins alpha (MIP-1α or CCL3; p=0.31) (**Fig. 4**). Collectively, the results suggest that NV1-based therapy could balance inflammatory responses and also could have anti-viral potency against SARS-CoV-2.

**Figure 3:**
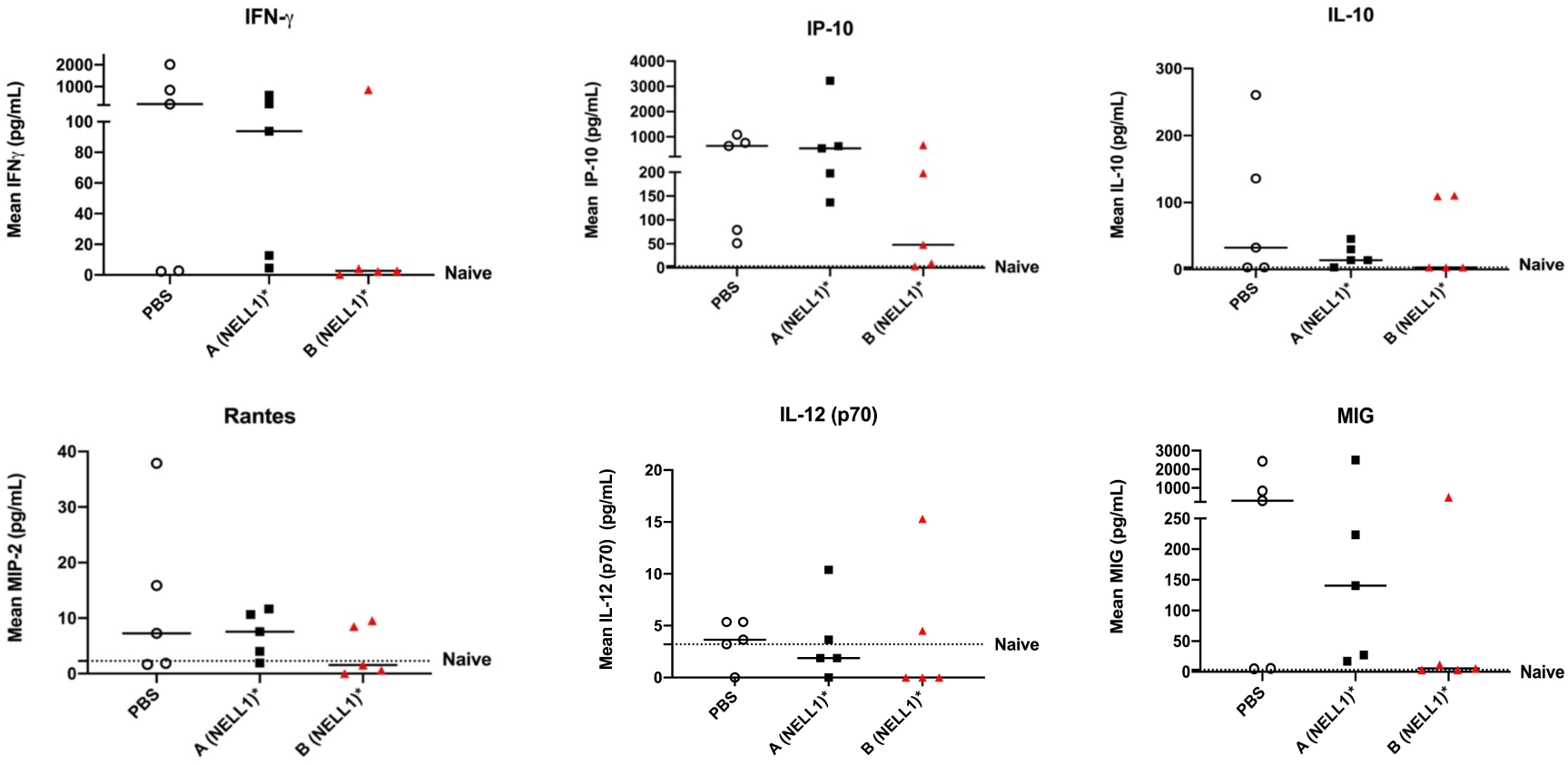
Inflammatory molecules that were decreased in BAL fluid samples from SARS-CoV-2 infected NV1-treated tg mice hACE2. The BAL fluid obtained from SARS-CoV-2-infected tg-mice hACE2 treated with NV1 either 1.25 mg/kg BW [A (NV1), N=5) or 2.5 mg/kg BW [B (NV1), N=5], and control (PBS-treated, N=5) was analyzed for cytokine and chemokine profile by Luminex assays. Relative levels of the cytokines, IFN-*γ*, IL-10, IL-12(p70), IP-10, MIG and Rantes, were decreased in one or both dosages are shown.

**Figure 4.**
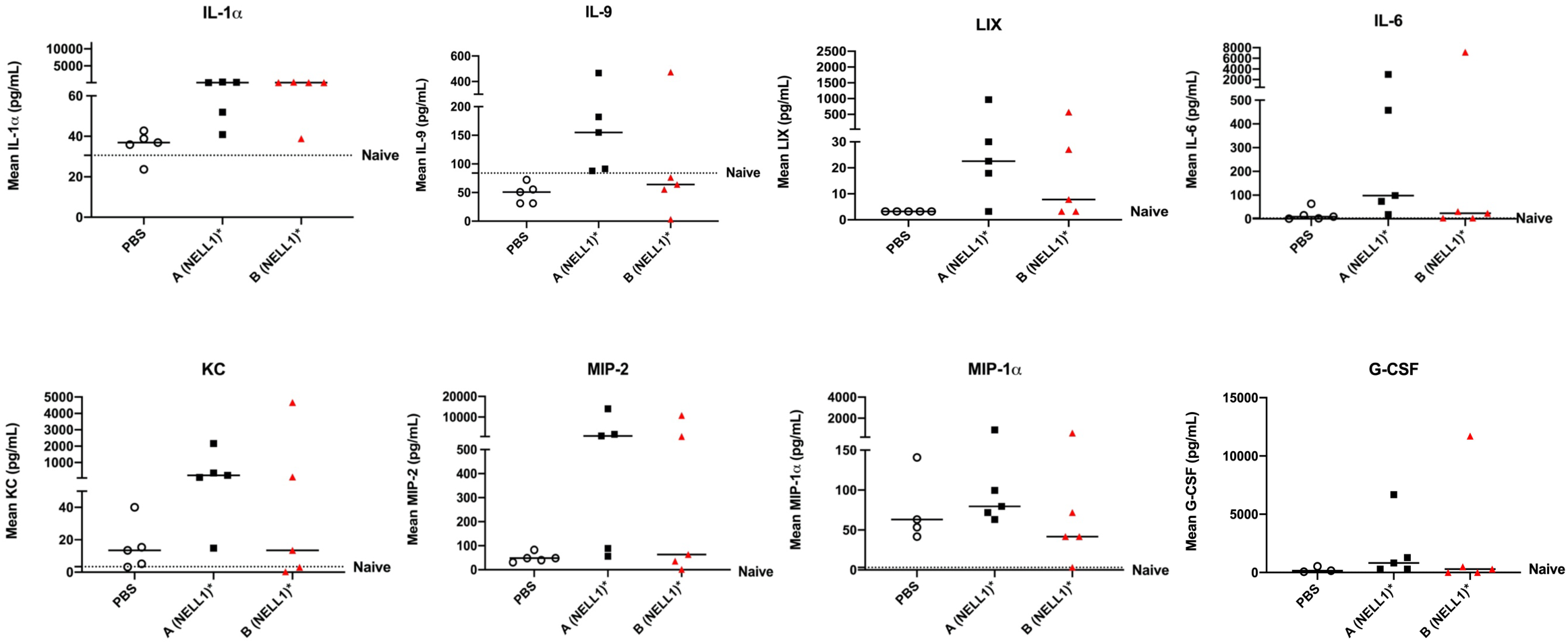
Inflammatory molecules that were increased in BAL fluid samples from SARS-Cov-2-infected NV1-treated tg-mice hACE2. The BAL fluid obtained from SARS-CoV-2-infected tg-mice hACE2 treated with NV1 either at 1.25 mg/kg BW [A (NV1), N=5) or 2.5 mg/kg BW [B (NV1), N=5], and control (PBS treated, N=5) was analyzed for cytokine and chemokine profile by Luminex assays. Relative levels of eight cytokines and chemokines (IL-1*α*, IL-9, IL-6, KC, LIX, MIP-2, MIP-1*α*, and G-CSF) were increased in one or both doses.

### SARS-Cov-2 infected hACE2 tg mice manifest pneumonia and vasculitis which are early signs of progression to severe COVID-19 in humans

Histopathological analyses of lung tissues from hACE2 tg mice infected with SARS-CoV-2 collected at time euthanasia identified common changes among many of the infected mice including: interstitial to broncho-interstitial pneumonia, increased circulating neutrophils (neutrophilic leukocytosis), occasional vasculitis, and perivascular edema (**Fig. 5**); occasional animals in each treatment group displayed no significant histologic findings. In affected mice, pulmonary inflammation was multifocal and affected 5-20% of the examined tissue and consisted primarily of infiltration of neutrophils and/or lymphocytes and macrophages into the alveolar septa and alveolar lumens. There were no detectable differences in pulmonary changes between treatment groups. There were no significant changes in any of the examined sections of lung from uninfected animals. The histologic characteristics are consistent with clinical observations of pulmonary damage in human COVID-19 patients that often accelerate to respiratory distress and critical disease if the viral load increases and standard of care treatments are ineffective.^6-7, 23^ The viral dose administered in this study was lower (2.8 × 10^3^ pfu) than the level used in previous published reports on this mouse model (2.8 × 10^4^ pfu), but resulted in the same rapid weight loss, behavioral and mortality phenotypes; hence, this treatment regimen is sufficient for generating clinically relevant disease features in hACE2 tg mice.^45^ This pilot study was designed to examine if NV1 treatment of SARS-CoV-2 infected hACE2 mice increased survival by mitigating hyperinflammation; hence, all control and treated mice were maintained until the euthanasia criteria were reached and lung tissues were collected at this point. As expected, no significant differences in pathologic features were observed among treatment groups (Data not shown). To address the direct effects of NV1 treatment on lung tissue healing, a follow-up study is underway to analyze lung tissues collected at the same timepoint after treatment (e.g. 24 hrs after last injection of NV1). The rapid body weight loss, survival profile, molecular and histopathology data obtained in this study all suggest the utility of the hACE2 tg mice as a model for rapid screening *in vivo* of novel therapeutics, like NELL1-NV1.

**Figure 5.**
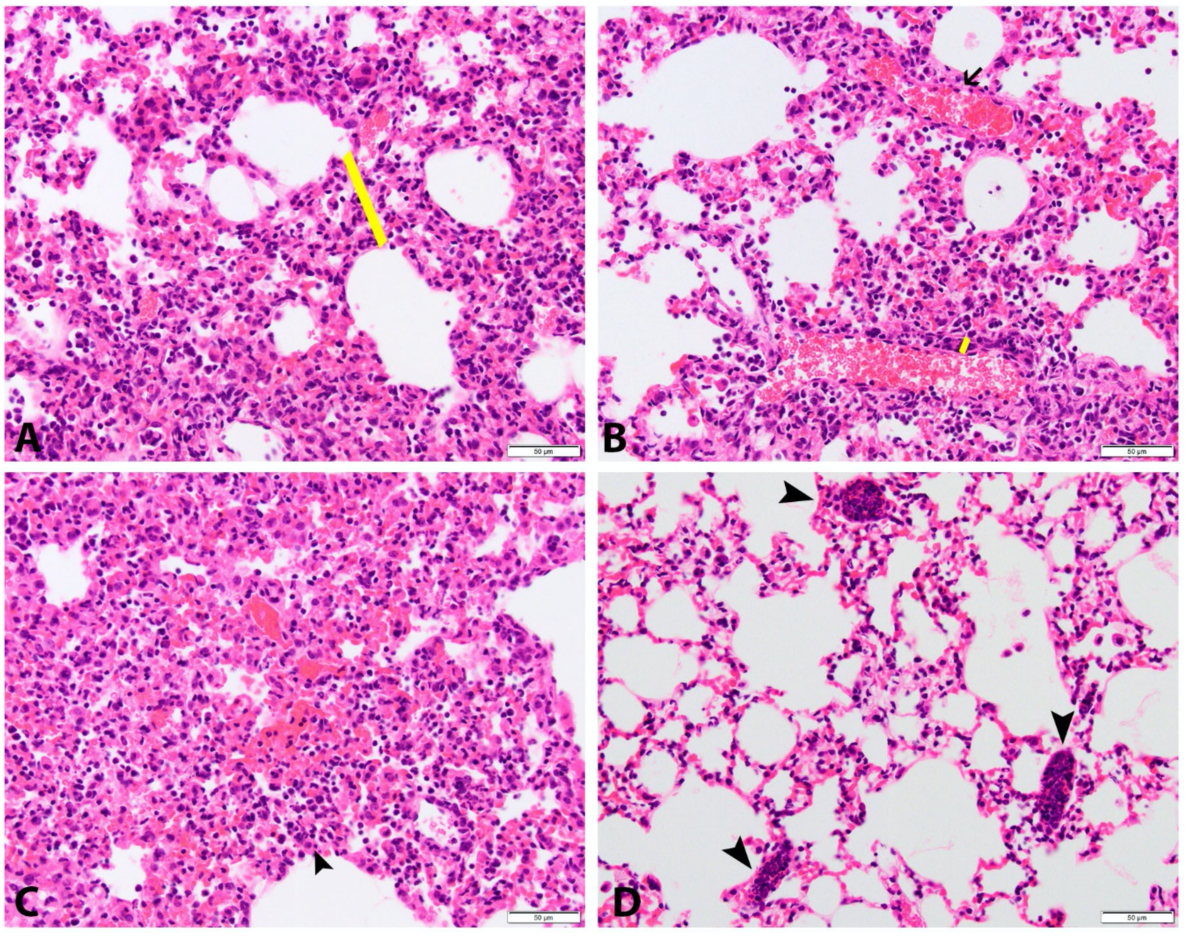
Histopathologic changes in lungs from SARS-CoV-2-infected tg-mice hACE2 harvested at euthanasia. Pulmonary changes included: **(A)**. expansion of the alveolar septa by neutrophils, lymphocytes, and macrophages (interstitial pneumonia), **(B)** expansion of the vascular wall (yellow bar) by neutrophils, karyorrhectic debris, and lymphocytes (vasculitis) compared to an unaffected vessel above (black arrow), **(C)** filling of alveoli by neutrophils, macrophages, and lymphocytes (black arrowhead), and **(D)** increased circulating neutrophils within capillaries (neutrophilic leukocytosis; black arrowheads). Hematoxylin and eosin, 40x magnification.

## DISCUSSION

The devastating impact of COVID-19 on public health, economy, medical personnel, healthcare infrastructure and sociocultural norms is rapidly accumulating as the COVID-19 pandemic extends into the winter season and the second year. Despite unprecedented efforts and coordination between public and private sectors, pharmacological products that safely and effectively slow disease progression and improve survival are not yet available.^1^ Despite the recent deployment of vaccines, there remains a need for effective therapeutics that (1) quell and/or balance the cytokine storm and the exacerbation of COVID-19 lung damage, and/or (2) promote lung tissue regeneration in COVID-19 survivors. In this pilot study, we tested for the first time, the efficacy of recombinant NELL1-NV1 protein as a therapy for COVID-19, using a murine model for COVID-19. We also evaluated if the pathways and mechanisms mediated by NELL1-NV1 in viral-induced tissue injury were similar to or overlapped with those observed in earlier studies of tissue healing and regeneration induced by the NELL1 protein.

There has been recent success in generating a small animal model of severe COVID-19 using the tg-mice hACE2 (under the control of the K18 promoter) to enable the understanding of the progression of viral infection from mild to life-threatening respiratory distress and death.^45-48^ The cytokeratin K18 promoter drives the high expression of the human ACE2 receptor in epithelial tissues that are primary targets of SARS-CoV-2 binding and entry in COVID-19. Several studies have now reported that this animal model recapitulates several features of the human disease.

With an intranasal delivery of 2.5 × 10^4^ plaque forming units (pfu) of SARS-CoV-2 (strain 2019n-CoV/USA_WAI/2020), Winkler *et al*. (2020) observed rapid weight loss beginning in 4 days and by 7 most animals lost 25% of body weight and appeared moribund.^45^ There were no sex differences in effects observed by Winkler et al. (2020). In contrast, Golden et al. (2020) female K18-hACE2 mice survived 60% higher than male mice at a lower viral dose (2×10^3^) of SARS-CoV-2, but not at 2.0×10^4^.^48^ This mouse model is therefore a valuable tool to test therapies to mitigate against severe manifestation of human COVID-19 and evaluate the potential of NELL1-NV1, a signaling protein that mediates healing after acute damage in various tissues. In this study, a 2.8 × 10^3^ pfu viral dose was delivered by intranasal injection and induced a dramatic weight loss in SARS-CoV-2 infected untreated control mice immediately at 1 dpi, deteriorated health condition (ruffled fur, hunched back and reduced mobility) and met criteria for euthanasia at 4-5 dpi for untreated infected tg-mice hACE2.

NELL1-NV1-treated animals at the lower dose of 1.25 mg/kg body weight administered at 0 and 3 dpi did not manifest significant weight loss until 6 dpi. The response of all females this treated group was particularly striking in that there was an increase in weight and were phenotypically normal for the first four days before weight loss began. The gender differences in the severity and mortality of COVID-19 is well documented since the early phase of the pandemic in certain patient cohorts and countries.^13,14^ More recently, a meta-analysis of global data consisting of 3,111,714 cases confirmed that males have 3 times the risk of severe COVID-19, requiring hospitalization, intensive care and higher odds of death compared to females.^49^ This disparity demands a closer examination in order to establish different clinical management strategies and protocols for the administration of therapeutics to specific patient groups. Since this pilot study used a small test group of females (N=3) and males (N=2) in the NV1-treated group showing a different pattern of BW loss between females and males, gender-based differences did not reach statistical significance. However, because of the importance of sex in COVID-19, this factor will be examined further in a follow-up study on a larger mouse population and using other doses of NV1. Future studies on NV1 in a larger animal population will confirm sex biased effects, explore a wider dose range (lower NV1 levels since the highest dose in this study was not efficacious as the lower dose) and different frequency and timing of administration can address these issues. It has been postulated that the variability in immune responses between sexes may explain these gender-based differences in COVID-19, thus therapeutics like NV1 that affects components of innate and adaptive immunity need to be administered differently to males and females to maximize efficacy.

The increased survival of NV1-treated tg-mice hACE2 was associated with multiple effects on the inflammatory profile. NV1 reduced the levels of several pro-inflammatory molecules: IFN-γ, IL-10, IL-12(p70), IP-10, MIG and Rantes. These are known biomarkers for severe COVID-19 cases and reveals a very pronounced NV1 effect in modulating the level of the inflammatory pathways mediated by interferon gamma.^15-18, 50-56^ IL-12 p(70) regulates the levels of IFN-γ, which in turns upregulates IP-10 and MIG.^53-56^ This is a tightly regulated pathway that is activated in response to the early phase of tissue injury and infection. Excessive activation of these molecules disrupts the T helper (Th1) cell-mediated immune responses that underlie many autoimmune diseases (arthritis, lupus, inflammatory muscle myopathies and severe reactions to viral and bacterial infections (HIV, RSV etc.).^57^ Interestingly, recent studies showed SARS-CoV-2-reactive IFN-γ CD8+ T cells in COVID-19 patients, and might be synergistic with IFN-α, which is currently being evaluated as a treatment in current COVID-19 clinical trials,^50,58^ MIG (monokine induced by IFN-γ), plays an important role in promoting inflammation and was correlated with disease severity in COVID-19 patients.^51,54, 59^ Rantes, a chemokine secreted by many cell types, promotes homing and migration of effector and memory T cells and viral clearance, is elevated in mild disease.^56, 60, 61^

Strikingly, numerous cytokines/chemokines were increased in NV1-treated mice, many of which are involved in viral clearance (e.g. IL-1α) and immune homeostasis (e.g. G-CSF).^62^Cytokines/chemokines involved in neutrophil recruitment, migration and activation were also increased, such as IL-9, LIX, MIP-2, KC and G-CSF, suggesting an important role of neutrophils in SARS-CoV-2 viral clearance and cytokine balance.^63-66^ Although there are numerous clinical reports on the increase or decrease of IL-6 in COVID-19, we observed an increased level in NV1-treated mice. There is no consensus on the levels and role of IL-6 as biomarker for disease progression but clinical trials using potent inhibitors of IL-6 show the increased propensity for secondary infections.^67^ This suggests that prolonged low levels of IL-6 are not desirable as it results in lowered immunity. Since the inflammatory profile in this study was obtained at euthanasia (not at the early onset of damage), it is possible that the higher level of IL-6 and other pro-inflammatory molecules is a positive effect of this therapy. Confirmation of this hypothesis will be tested in follow-up studies examining the NV1 inflammatory effects at earlier timepoints and the cytokine/chemokine levels in lung and peripheral blood/serum (in addition to BAL).

Previous studies have shown that NELL1 and NV1 proteins regulates the inflammatory response to promote tissue healing after acute injuries and in systemic inflammatory diseases (**Fig. 1**; e.g. wounds, bone fractures and osteoarthritis). In this study, NV1 effects on the levels of IL-*β*, IL-8, TNF-*α* and the transcription factor NF-*κ*B were not detected in the cytokine profile of BAL from SARS-CoV-2 infected mice. Since cytokine analysis was conducted at euthanasia when all mice were exhibiting severe symptoms instead of a fixed time post-administration of NV1, it is possible that the established effects of NELL1/NV1 are exhibited early after treatment. Alternatively, the impact of NELL1 in viral induced tissue damage may involve modulating different pathways in the inflammation cascade than in acute tissue injuries and other inflammatory diseases.

Recent immune profiling and genomics analyses of severe COVID-19 patients show that a variety of molecules are predictive markers and targets for possible therapeutic strategies.^16, 55, 68^ The immune profiles of patients who recovered from moderate COVID-19 were found to be enriched in tissue reparative growth factor signature A, whereas the profiles of those with who developed severe disease had elevated levels of other molecular signatures. Signature A contained several stromal growth factors, including epidermal growth factor (EGF), platelet-derived growth factor (PDGF) and vascular endothelial growth factor (VEGF), that are systemic mediators of wound healing and tissue repair.^16^ The NELL1 protein’s efficacy in wound healing and tissue repair suggests that it may activate similar processes in restoring tissue damaged by SARS-CoV-2 infection.

In recent global human clinical trials (405 sites in 30 countries, 11,266 patients), the top four candidates hydroxychloroquine, remdesivir, lopinavir and interferon-β1a, which were strongly inhibitors of pro-inflammatory cytokines, viral infection and replication did not significantly reduce mortality, initiation of ventilation or hospitalization for severe COVID-19.^28, 69^ We propose that a therapy such as NELL1-NV1 which mitigates the cytokine storm by modulating multiple cytokines and balancing the levels of inflammatory molecules necessary for an effective immune response against infection will establish balance in the aberrant immune response to SARS-CoV-2 infection.^74^ These biological effects are likely to promote survival and recovery from lung tissue damage.^65, 75^ Further testing of this hypothesis will be conducted in larger statistically powered study with a wider escalating dose range (lower doses than 1.25 mg/kg), histopathology of lung tissue to examine the severity of tissue injury and scarring, analysis of immune cell populations and examining early (after first administration of NV1) and late effects (at euthanasia) on the levels of the viral load and inflammatory molecules.

## FOOTNOTES

### Disclaimer

The views expressed are those of the authors and do not reflect the official policy or position of the Uniformed Services University of the Health Sciences, the Department of Defense, or the United States government.

## Abbreviations used

COVID-19,: coronavirus disease-19;
SARS-CoV-2,: severe acute respiratory syndrome coronavirus 2;
hACE2,: human angiotensin I-converting enzyme 2 receptor; neural epidermal growth factor like 1 protein variant (NV1)

## Acknowledgments

This study was supported by USUHS intramural grant [RB] and NellOne Therapeutics Inc. private funds [CTC, SE]. Maciel Porto monitored lab processes and performed Luminex assays, Melissa Hamilton analyzed the data, and Megan Lok managed the *in vivo* part of the study (Bioqual Inc.). Anwar E. Ahmed at the department of Preventive Medicine and Biostatistics (USUHS, Bethesda, MD) conducted the statistical analysis for this study.

